# Single-cell analysis of progenitor cell dynamics and lineage specification of the human fetal kidney

**DOI:** 10.1101/258798

**Authors:** Rajasree Menon, Edgar A. Otto, Austin Kokoruda, Jian Zhou, Zidong Zhang, Euisik Yoon, Yu-Chih Chen, Olga Troyanscaya, Jason R. Spence, Matthias Kretzler, Cristina Cebrián

**Affiliations:** Department of Computational Medicine and Bioinformatics, University of Michigan, Ann Arbor; Department of Internal Medicine, Division of Nephrology, University of Michigan, Ann Arbor; Department of Internal Medicine, Division of Gastroenterology, University of Michigan, Ann Arbor; Department of Cell and Developmental Biology, Division of Gastroenterology, University of Michigan, Ann Arbor; Lewis-Sigler Institute for Integrative Genomics, Princeton University, Princeton, New Jersey; Graduate Program in Quantitative and Computational Biology, Princeton University, Princeton, New Jersey; Department of Electrical Engineering and Computer Science, Department of Biomedical Engineering, University of Michigan, Ann Arbor; Flatiron Institute, Simons Foundation, New York, New York; Department of Computer Science, Princeton University, Princeton, New Jersey

## Abstract

The mammalian kidney develops through repetitive and reciprocal interactions between the ureteric bud and the metanephric mesenchyme to give rise to the entire collecting system and the nephrons, respectively. Most of our knowledge of the developmental regulators driving this process has been gained from the study of gene expression and functional genetics in mice and other animal models. In order to shed light on human kidney development, we have used singlecell transcriptomics to characterize gene expression in different cell population, and to study individual cell dynamics and lineage trajectories during development. Single cell transcriptome analyses of 3,865 cells identified 17 clusters of specific cell types as defined by their gene expression profile, including markers of ureteric bud tip- and metanephric mesenchyme-specific progenitors, as well as their intermediate and differentiated lineages including the mature collecting ducts, the renal vesicle and comma- and s-shaped bodies, immature and mature podocytes, proximal tubules, loops of Henle and distal tubules. Other lineages identified include mesangium and cortical and medullary interstitium, endothelial and immune cells as well as hematopoietic cells. Novel markers for these cell types were revealed in the analysis as well as components of key signaling pathways driving renal development in animal models. Altogether, we provide a comprehensive and dynamic gene expression array of the human developing kidney at the single-cell level.

## INTRODUCTION

The development of the embryonic kidney is a paradigm for the complex inductive and regulatory mechanisms driving organogenesis (Grobstein, 1953). In mammals, the epithelial ureteric bud (UB) tips undergo repetitive branching morphogenesis until birth and generate the entire collecting system including the ureter, calyces and collecting ducts but not the bladder (Chi et al., 2009; Riccio et al., 2016; Shakya et al., 2005). Around the tips of the UB, nephron progenitors condense into cap mesenchyme, epithelialize and undergo complex morphogenesis to differentiate into the vast majority of cells in the nephron including the epithelial cells of the Bowman’s capsule, the podocytes, proximal and distal tubules, loops of Henle and the connecting tubule (Cebrian et al., 2014; Kobayashi et al., 2008; Mugford et al., 2008). Interspersed between the nephron progenitors, the interstitium promotes survival and differentiation of the progenitors as well as branching of the UB and contribute to the mesangial and endothelial lineage (Das et al., 2013; Mugford et al., 2008).

Studies in mice have identified several signaling pathways that are cornerstone of the reciprocal epithelial-mesenchymal interactions driving kidney development. For example, the soluble ligand Glial cell-line Derived Neurotrophic Factor (Gdnf) is expressed by the mesenchymal nephron progenitors and signals through the epithelial UB tip-specific receptor tyrosine kinase Ret, the co-receptor Gfra1 and their downstream transcription factors Etv4 and Etv5. This signaling cascade is critical for UB outgrowth and branching (Cacalano et al., 1998; Lu et al., 2009; Pachnis et al., 1993; Pichel et al., 1996; Schuchardt et al., 1994). Wnt9a expressed by the stalk of the UB induces Wnt4 expression in the early nephron, and canonical wnt signaling regulates the balance between self-renewal and differentiation of nephron progenitors (Carroll et al., 2005; Park et al., 2007). On the other hand, Wnt5a mediates non-canonical wnt signaling that ensures proper positioning of the Metanephric Mesenchyme (MM) and UB outgrowth (Huang et al., 2014; Pietila et al., 2016; Yun and Perantoni, 2017). Fibroblast Growth Factors (Fgfs) and their receptors are also required for kidney development. Fgfr2 is required for UB branching (Zhao et al., 2004), Fgf9 and Fgf20 regulate nephron progenitor self-renewal (Barak et al., 2012) and Fgf8 is required for progression of nephrogenesis from the renal vesicle (Grieshammer et al., 2005; Perantoni et al., 2005). In addition, Notch signaling promotes further differentiation into proximal and distal segments of the nephron (Cheng et al., 2007; Surendran et al., 2010), directs differentiation of nephron progenitors (Boyle et al., 2011) and drives specification of stromal cells into mesangium (Boyle et al., 2014).

Our knowledge of the lineage relationships between the renal cell types and these regulatory pathways driving renal morphogenesis and differentiation arise primarily from the study of mouse models. While the murine and the human kidney seem to share a common developmental pattern, they also present anatomical, physiological and pathophysiological differences, as well as significant differences in gene expression (O'Brien et al., 2016), suggesting that significant new information can be gained by directly studying human kidney development.

The complexity of the developing human kidney, with over a dozen different mature cell types and in the process of differentiating in parallel, has precluded a systematic analysis of its expression profile. In this context, single-cell transcriptomics provides a unique tool to analyze individual cell types from progenitors to differentiated cells as well as their intermediate transitional states.

We have used DropSeq to perform single cell sequencing on 3,865 cells from 3 developing human kidneys and identified 17 distinct cell populations based on their expression profile. These populations include progenitors, immature and mature renal cell types and proliferating cells. We further identify new markers for specific cell types in the developing human kidney, and we use computational approaches to infer cell lineage trajectories and interrogate the complex network of signaling pathways and cellular transitions during development.

## RESULTS AND DISCUSSION

### Single cell sequencing identifies progenitors, mature and intermediate cell types in the developing human kidney

Analysis of single cell sequencing data of 3,865 cells from 3 human embryonic kidneys revealed 19 clusters. Using cell clustering (Fig. 1B-C and Sup. Fig.1A) and lineage trajectory algorithms (Fig. 1D), we have assigned cell identity to 17 of these cell clusters. Using pre-defined anchor genes cells were assigned an identity based on expression of genes in single cells (Fig. 1C, Sup. Table 1) that are predicted to be expressed in specific cell types based on prior literature (Table 1). We have also performed lineage trajectory analysis, which uses an unbiased computational approach to infer lineage relationships to distinguish between progenitor cell types (Fig. 1D). Both analyses demonstrate that differentiated cell types are located in the periphery of the cluster plot (Fig. 1B) and trajectory plot (Fig. 1D) while progenitor cells lie inside the plots. Based on these analyses, the identities of the 17 clusters are as follows: Clusters 14,15 and 17 are well-defined, compact and isolated clusters that correspond to endothelial, immune and red blood cells respectively. Also in the periphery but not as neatly isolated are clusters 5 and 10 that correspond to cortical and medullary interstitial cells respectively; the more mature medullary interstitium is located towards the periphery. Clusters 9 and 13 also correspond to the immature and more mature podocyte populations respectively. Cluster 8 includes loops of Henle as well as distal tubule cells. Clusters 4 and 18 correspond to the developing and mature collecting duct epithelia respectively and cluster 6 includes proximal tubule cells. The center of the cluster plot is also populated by progenitor and proliferating cells. The cap mesenchyme that contains the nephron progenitor cells belongs to cluster 1 while cluster 2 is a less compacted cluster that corresponds to the intermediate stages, ranging from pre-tubular aggregate to s-shaped body. This cluster overlaps to some extent with neighboring clusters for the more mature cell types like proximal, distal, Henle’s loop and even immature podocytes, confirming their lineage relationship. Proliferating cells cluster in 3 different groups: 3, 7 and 11. While cells in all three of these clusters express proliferation genes like *MKI67*, they likely represent different stages of cell division; *CKD1* controls cell division during mitosis and is expressed by cells in cluster 7, while cells in cluster 11 express *CKD2* and *CKD4* that are expressed during the S/G2 phase of cell division. Most interestingly, cells in cluster 16 express UB tip specific genes but sit directly adjacent to the proliferating cluster 3 rather than the mature and immature collecting duct clusters (18 and 4 respectively). Cluster location, together with the expression of proliferating genes, suggests that cells in cluster 16 correspond to proliferative UB tip cells. Cluster 0 has not been assigned an identity and includes those cells that express housekeeping genes but don’t express enough cell-type specific genes. These cells could be damaged or have lower reads during sequencing and not enough information was retrieved to assign them to a cluster. Similarly, cluster 12 has not been assigned an identity and is a relatively loose cluster positioned between the cap mesenchyme cells, proliferating cells and developing epithelia; while they likely represent an intermediate population, the small number and power of the highly variable genes identified in this cluster (Sup. Table 1) precludes assigning a specific identity with confidence. All together, our approach identifies cell populations that are consistent with the major renal mature cell types, their progenitors, as well as intermediate stages of differentiation.

**Figure 1.**
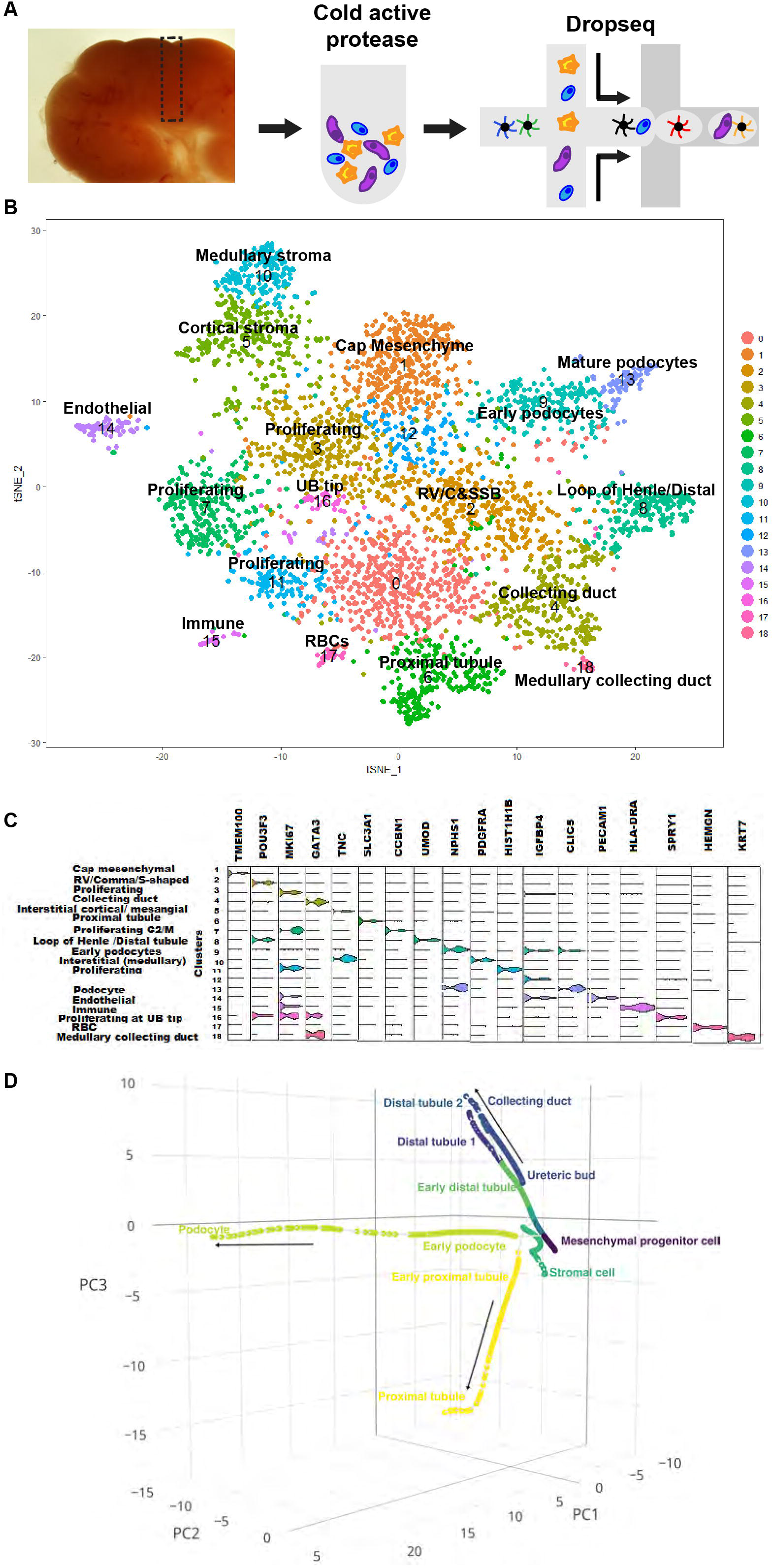
Single cell transcriptomic and trajectory analysis identifies progenitor and differentiated cells in the developing human kidney. A: Diagram representing the processing of embryonic samples for dropseq. Small kidney fragments encompassing from the cortex to the inner medulla are excised from the sample, digested and subjected to microfluidic co-flow with barcoded beads. B: t-SNE cluster map and the identity assigned to each cluster based on their gene expression profile. Each dot in the plot represents a single cell. C: Violin plot of representative genes for each cluster. D: Trajectory analysis of single-cell RNA-seq data. Major mature cell types and their progenitors are labeled in the plot. Each dot in the plot represents a single cell, shown with its position on the estimated developmental trajectory. The dots are connected by lines, which indicate the estimated trajectories. The direction of differentiation is indicated by arrow. The trajectories in transcriptome space are visualized in the space spanning the first three principle components of the single RNA-seq data.

**Table 1:**
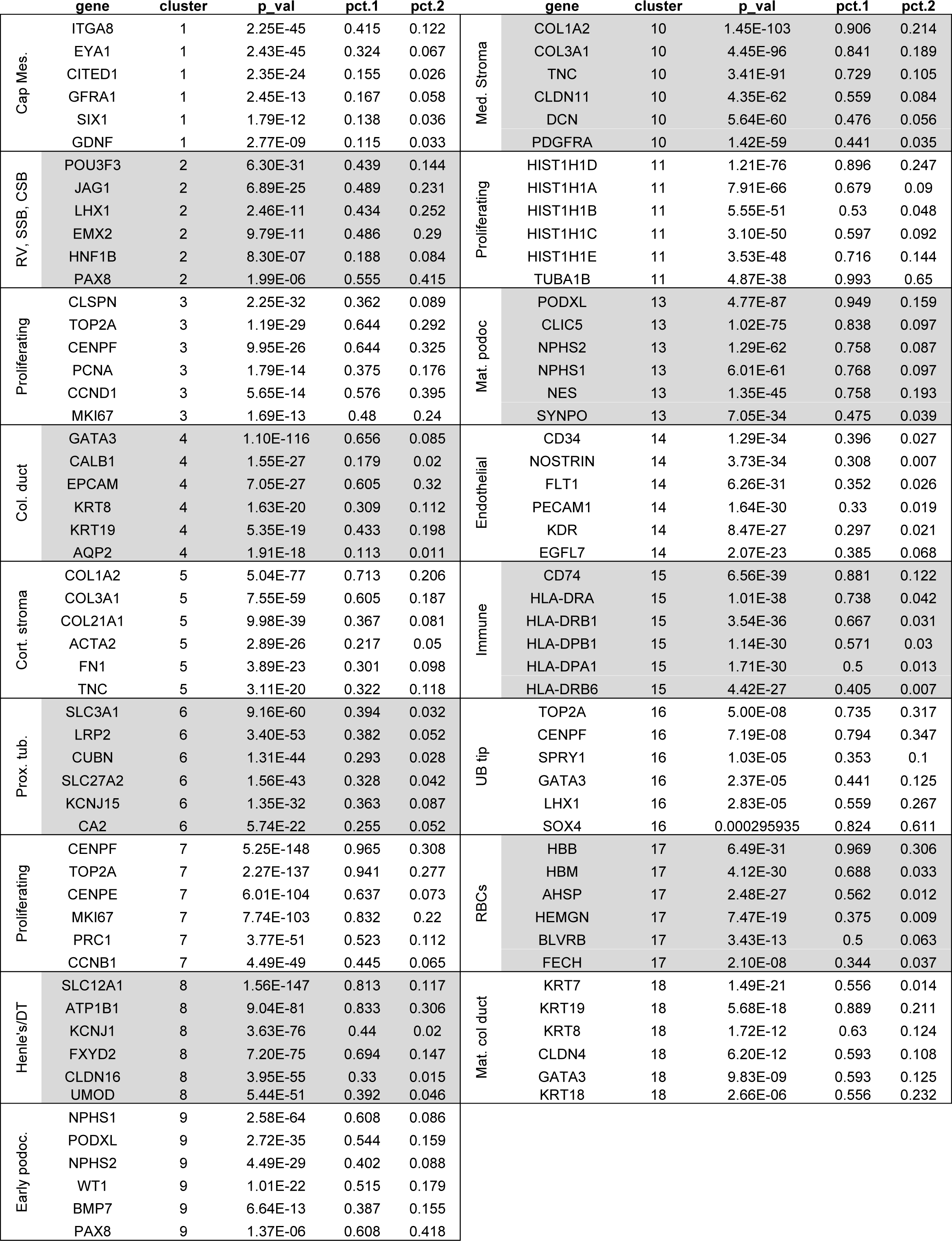
Representative genes for each cluster: Pct1: Percentage of cells in the cluster of interest. Pct2: Percentage of cells in all other clusters.

### Analysis of cluster-specific gene expression identifies novel marker genes

The identity of each cluster is assigned based on known gene expression patterns either in human or in mouse developing kidneys (Table 1); however, new markers for these clusters are also revealed by the analysis. Cluster 1 corresponds to the nephrogenic mesenchyme and is characterized by the expression of *ITGA8, EYA1* and *TMEM100* among other genes (Muller et al., 1997; Rumballe et al., 2011; Xu et al., 1999 and sup. fig. 1). Cluster 1 also expressed two genes that have not been previously associated with nephrogenic mesenchyme, *ELAVL4*(ELAV like RNA binding protein 4) and *ECEL1* (endothelin converting enzyme like 1) (Sup. Fig. 2). In addition, we detected strong expression of *COL2A1*, which is a well-established chondrocyte marker (Ng et al., 1993) and *ETV1* (ETS translocation variant 1) in the nephrogenic mesenchyme. The expression of *ETV1* in the developing human kidney is in contrast with previous data from embryonic mouse kidney where *Etv1* expression was not detected (Lu et al., 2009). COL2A1 is expressed in the developing mouse kidney (Wada et al., 1997) but its cell specificity and functional role remain to be elucidated. The specific expression of these two genes in the nephrogenic mesenchyme of human developing kidneys suggests a possible novel role in nephron induction and/or differentiation. In the collecting duct (Cluster 4), the genes most significantly defining the cluster were *GATA3* and Branched Chain Aminotransferase 1 *(BCAT1)*(Sup. Table 1). While *BCAT1* has not been studied in the kidney nor its expression reported, it has been recently shown to regulate early liver bud growth in the developing mice and in human embryonic stem cell derived liver organoids (Koike et al., 2017). Specific to the immature podocytes (Cluster 9) we find the expression of Olfactomedin 3 *(OLFM3)* (Fig. 3C). The function of OLFM3 has not been elucidated but it has been suggested to be involved in cell adhesion (Hillier and Vacquier, 2003). On the other hand, mature podocytes (Cluster 13) show specific expression of Netrin G1 *(NTNG1)* (Sup. Table 1), a member of the Netrin family. Netrins are extracellular, laminin-related proteins that provide guidance for migrating cells and NTNG1 is a membrane-tethered glycophosphatidylinositol (GPI)-linked Netrin (Lai Wing Sun et al., 2011). Expression of *NTNG1* has been reported in the adult human kidney by Northern blot and semiquantitative PCR (Meerabux et al., 2005) but its specific podocyte expression in the developing kidney has not been reported. These are just a few examples of the novel cluster-specific genes identified by our single-cell transcriptomics analysis, that open new avenues to further our understanding of human kidney development. Additional studies will be required to characterize the role of these expressed genes, and to determine their functional relevance, possibly through the use of cultured human tissue, or via human renal organogenesis.

**Figure 3:**
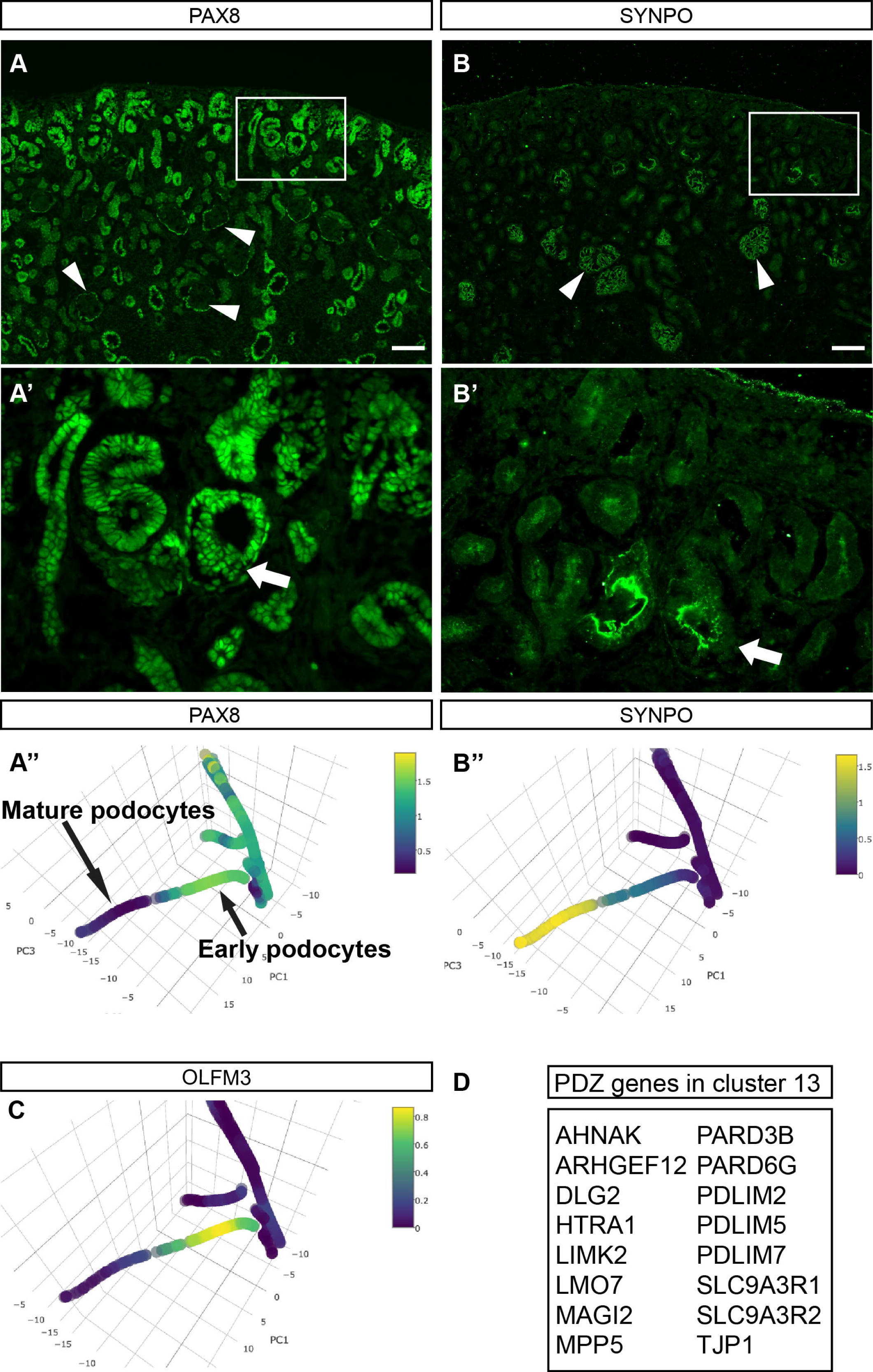
Immature and mature podocytes can be identified by their gene expression profile. A, A’ and A’’: Immunofluorescence and trajectory analysis detecting PAX8 expression. Arrowheads in A point at mature glomeruli where PAX8 is expressed in the epithelial cells of the Bowman’s capsule. Arrow in A’ points at an immature glomerulus where PAX8 is expressed in both the epithelial cells of the Bowman’s capsule and the developing podocytes. B, B’ and B’’: Immunofluorescence and trajectory analysis detecting SYNPO expression. Scale bar 100um.

### Specification of renal cell lineages is conserved in the human kidney

Conventional genetic lineage tracing cannot be performed on human tissues for obvious reasons. In mice, genetic studies have identified the tips of the UB as the progenitors of the entire renal collecting system (Chi et al., 2009; Riccio et al., 2016; Shakya et al., 2005) and the cap mesenchyme as the progenitor for the nephron epithelia (Cebrian et al., 2014; Kobayashi et al., 2008; Mugford et al., 2008). In a similar fashion, the cortical stroma contains the progenitors of the renal stroma although some endothelial cells arise from extra-renal contribution (Das et al., 2013). Our results from cluster analysis and from trajectory analysis indicate the transition in gene expression from progenitor populations to differentiated ones (Fig. 1D). This analysis identifies 5 different cellular trajectories. The UB lineage is separated from the mesenchyme-podocyte and mesenchyme-tubular segments, reflecting our understanding of the developmental origins of these tissues in mice. Proximal and distal tubules as well as the podocyte trajectory originate from the mesenchymal progenitor cell segment. The stromal or interstitial population is also separate from the UB and the mesenchymal lineages but it is in close proximity to the latter reflecting the overlapping expression of genes common to both mesenchymal progenitors and stroma, such as *ITGA8, NR2F1* and *NR2F2.* In most cases, progenitor and differentiated clusters remain together, with undifferentiated populations positioned more central in the tSNE plot (Fig. 1B). Three very clear examples are the interstitial cells, the podocytes and the collecting system. In the case of interstitial cells, cluster 10 contains the differentiated, medullary interstitium characterized by the expression of MCAM, Tenascin (TNC) and Decorin (DCN) (Fig. 2A, C and E). The cortical interstitial cells in cluster 5 on the other hand have strong ACTA2, MCAM and TNC expression (Fig. 2A, B and E). In mice, MCAM is critical for kidney vasculature development and it is expressed in interstitial cells that differentiate into PECAM-expressing endothelial cells (Halt et al., 2016). Interestingly, several MCAM expressing cells are located in the cortical stroma (cluster 5) as well as in the endothelial cluster (cluster 14) (Fig. 2E and F), supporting the notion that MCAM-expressing cells differentiate into PECAM-expressing cells in the human developing kidney. Collagens are also prominently expressed in interstitial cells and both cluster 5 and cluster 10 express a plethora of collagen genes (Fig. 2D). Some collagens are more prominently expressed in each of the clusters; specifically, cells in cluster 5 express *COL12A1* while cells in cluster 10 express *COL5A1, COL7A1, COL16A1, COL25A1* and *COL27A1.* Hence, our data suggests that the human interstitium transitions from a *COL12A1 +, ACTA2+, MCAM+* positive immature state to a Decorin positive, high Tenascin mature interstitium and *PECAM+* endothelium.

**Figure 2.**
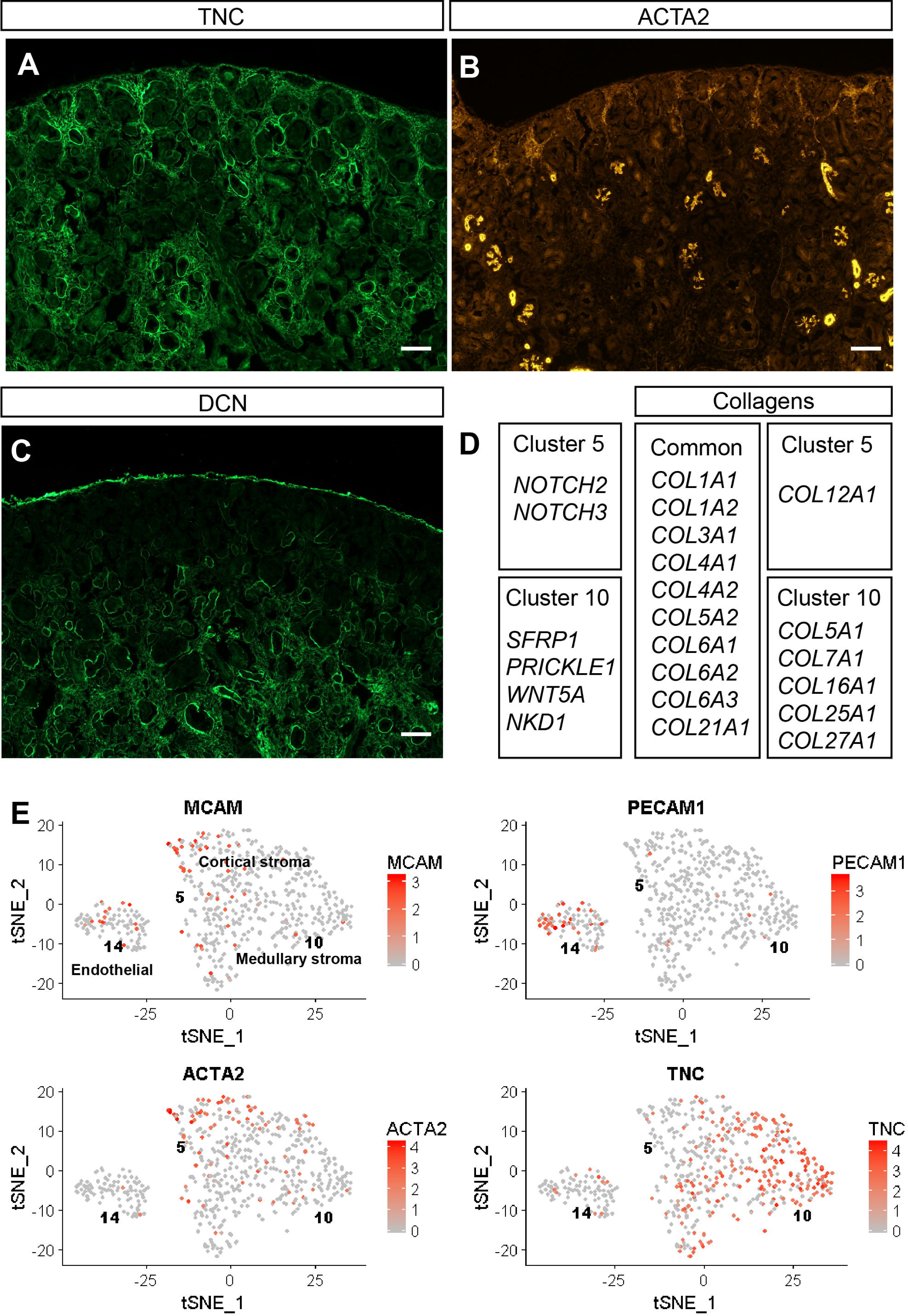
Human renal interstitium is divided into cortical and medullary clusters. A: Tenascin C protein is detected by immunofluorescence in both cortical and medullary areas of the human embryonic kidney. B: aSMA is present in the cortical stroma of the human developing kidney, in the glomerular mesangial cells and in perivascular cells. C: Immunofluorescence assay showing decori expression exclusively in the medullary interstitium. D: List of collagens and signaling molecules with differential expression between cortical and medullary interstitium. E: Subplots of endothelial and cortical and medullary interstitium. With a few exceptions PECAM1 expression is exclusive to the endothelial cluster while ACTA2 and Tenascin C are specific for the stroma. MCAM, a marker of developing endothelial cells is expressed in the cortical stromal cells as well as the endothelial cells. Each dot in the plot represents a single cell. Scale bar 100um.

In a similar fashion podocytes can be clustered into two cell populations, an immature one in cluster 9 characterized by the expression of *PAX8, BMP7, OLFM3* and *DSC2* and a mature population in cluster 13 that expresses high *Synaptopodin (SYNPO)* and *Dendrin* (DDN). Both clusters express known markers for podocytes including *PODXL*, Nephrin *(NPHS1)*, Podocin *(NPHS2)*, Nestin *(NES)* and *MAGI2* among others. In cluster 9, *PAX8* (Fig. 3A, A’ and A’’), *BMP7* and *DSC2* have been previously reported to stain immature podocytes (Garrod and Fleming, 1990; Kazama et al., 2008; Ohse et al., 2009) and *OLFM3* (Fig. 3C) is a new marker of developing podocytes. Exclusively in the mature podocyte cluster 13 we find podocyte-specific genes *Dendrin (DDN)* and *PLA2R1* (Lindenmeyer et al., 2010; Patrakka et al., 2007) and the novel marker NetrinG1 *(NTNG1).* A number of PDZ domain proteins are expressed in the mature podocyte cluster (Fig. 3D). Cluster 13 enrichment in PDZ domain proteins is consistent with a role for these proteins in establishing cell-cell contacts and the slit diaphragm characteristic of mature podocytes. Indeed, *MAGI2, PDLIM2, MPP5, PARD3B* and *TJP1* are associated with cytoskeletal and barrier function in podocytes (Ihara et al., 2014; Itoh et al.,2014; Koehler et al., 2016; Lu et al., 2017; Sistani et al., 2011). Our data suggest that several other PDZ domain proteins play a role in the establishment or maintenance of the slit diaphragm in mature human podocytes. This clear distinction between immature and mature podocytes could reflect a transition between pre-functional and functional filtering structures and the identification of novel markers for each of those populations provides additional tools in the study of podocyte function during development, in homeostasis and during renal disease.

Immature and mature collecting duct cells (clusters 4 and 18 respectively) sit adjacent to each other, with mature medullary cells expressing *KRT7* and *TACSTD2* (Tsukahara et al., 2011) and less mature cortical collecting duct cells expressing *GATA3* and *CALB1* (Fig. 1C). The UB tip cells (cluster 16), progenitors of both immature and mature collecting ducts (clusters 4 and 18 respectively), do not cluster next to their progeny but away from it in between the proliferating cells clusters (Fig. 1B). This likely reflects the highly proliferative nature of these cells, and the possibility that the transcriptome of proliferative cells differs significantly from that of their progeny. Indeed, a correlation plot for clusters 3, 4, 16 and 18 reveals the proximity of the UB tip cells to the proliferating cells (Sup. Fig. 1B). However, the UB tip cells appear in a continuum with the mature collecting duct cells in the trajectory analysis confirming their lineage relationship. The UB tip cluster cells express tip specific genes like *SPRY1, SOX4* and *ARL4C*(Huang et al., 2013; Matsumoto et al., 2014; Zhang et al., 2001) but also proliferation specific genes like *TOP2A, CENPF* and *BUB1B.*

### Pathways relevant to kidney development are present in the developing human kidney

Several signaling pathways drive renal organogenesis. Among them, Gdnf signaling via the receptor tyrosine kinase Ret and the co-receptor Gfra1 is critical for kidney development. Gdnf is expressed in the metanephric mesenchyme while UB tip cells express Ret and Gfra1 is expressed in both compartments (Ehrenfels et al., 1999; Golden et al., 1999; Pachnis et al., 1993). We identify *GDNF* and *GFRA1* expression in cluster 1 that contains the nephrogenic mesenchyme and *GDNF* expression is also shown in the progenitor segment of the trajectory analysis (Fig. 4C); *RET* is present in the collecting duct progenitors in the trajectory analysis as well as by immunofluorescence with RET specific antibodies on human embryonic kidney sections (Fig.4A and A’). RET is detected in the membrane of the cells at the tips of the branching UB, with some expression extending towards the trunk. Etv4 and Etv5 are downstream targets of Gdnf/Ret signaling (Lu et al., 2009) and drive UB branching in mice (Kuure et al., 2010; Lu et al., 2009; Riccio et al., 2016). They are expressed at the tip of the UB and the developing nephrons. In the human developing kidney we can detect ETV4 transcription factor in the nucleus of the tip cells as well as in the early renal vesicles (Fig.4B) and both ETV4 and ETV5 are detected in the UB tips and early renal tubules in the trajectory analysis (Fig.4B’ and D). Hence, the presence in the human developing kidney of these genes critical for this signaling pathway, suggests that this pathway is likely critical for human kidney development.

**Figure 4:**
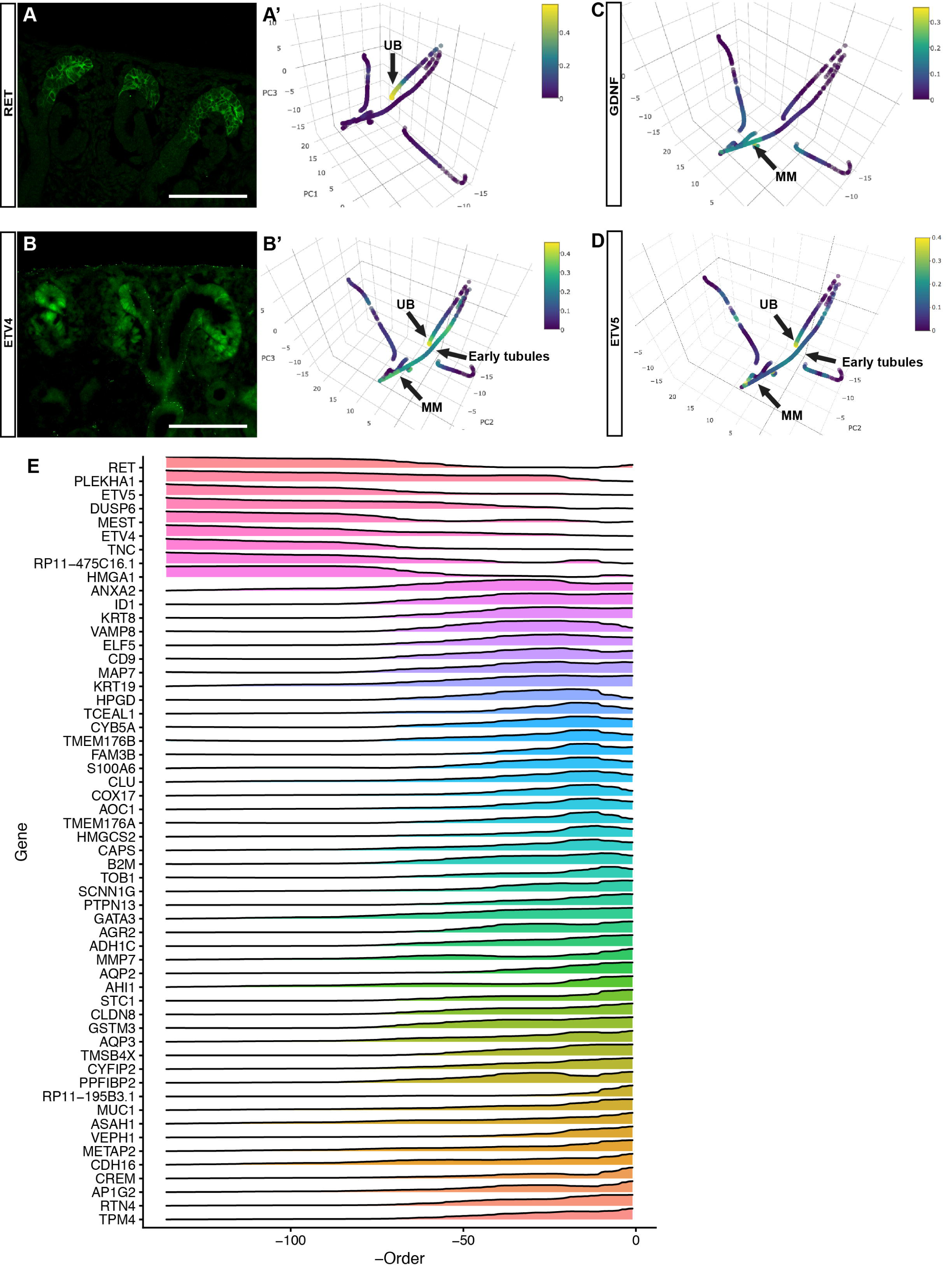
Gene expression in the human collecting duct. A, A’: Expression of the receptor tyrosine kinase RET is detected at the tips of the human ureteric buds by immunofluorescence and by trajectory analysis. B, B’: Immunofluorescence and trajectory analysis of ETV4, a downstream target of RET/GDNF signaling, that is also detected at the tips of the ureteric bud and in the developing renal tubules. C: Trajectory analysis of GDNF shows expression in the metanephric mesenchyme. D: Expression of ETV5 is detected in the UB tips and early nephrons by trajectory analysis. E: Gene expression heatmap for the collecting duct differentiation. Note RET expression in immature collecting duct cells (UB tip cells) and AQP2 expression in mature cells. Scale bar 100um.

The Notch signaling pathway is required for the differentiation of stromal cells into mesangium (Boyle et al., 2014), and both *NOTCH2* and *NOTCH3* are expressed by cells in the cortical interstitial cluster 5. Cells in this cluster express the mesangial cell markers *ACTA2, LGALS1*and *PDGFRB* suggesting that cluster 5 is composed of immature interstitial and mesangial cells. On the other hand, cells in cluster 10, the medullary interstitium, express *SFRP1, PRICKLE1, WNT5A* and *NKD1* suggesting that non-canonical WNT activity in the medullary interstitium drives planar cell polarity in neighboring epithelial cells. Surprisingly, *WNT5A* is the only *WNT*gene with significant expression in a single cluster (Sup. Fig.3). *WNT4, WNT7B* and *WNT11* are present in a few cells but are not enriched in any given cluster. A similar expression pattern is observed for *FGFs* and their receptors. Both *FGFR1* and *FGFR2* are expressed in the developing human kidney, with stronger expression and wider distribution for *FGFR1* (Sup. fig.4) and without evident enrichment in any cluster. *FGF7* shows clear enrichment in medullary and cortical stroma, in accordance with its reported expression in the developing kidney (Mason et al., 1994), however very little expression is detected for *FGF8-10.* While the low expression of many of these genes may represent a divergence between mouse and human development, we cannot preclude the possibility that their expression is too low to be detected or that they escape detection with the methods used.

## MATERIALS AND METHODS

### Single cell dissociation of human fetal kidney tissue using a cold active protease (Subtilisin)

All research utilizing human fetal tissue was approved by the University of Michigan institutional review board. Normal human fetal kidneys were obtained from the University of Washington Laboratory of Developmental Biology. All tissues were shipped overnight in Belzer’s solution at 4 degrees Celsius and were processed immediately upon arrival to the laboratory. Single cell dissociation was performed using a cold active protease as described in Adam et al. (Adam et al., 2017). One kidney was dissected in ice-cold PBS and finely minced in a petri dish on ice using razor blades. About 20 mg of tissue were added to 1 ml of cold active protease solution (PBS, 10 mg of *Bacillus Licheniformis* protease [Sigma, #P5380], 5 mM CaCl_2_, 20 U DNAse I [Roche, #4716728001]). The tissue was incubated in a 2 ml reaction tube for 15-20 min on a slow moving shaker (nutator) in a coldroom at 4°C with repeated trituration steps for 20 seconds every 5 minutes. Single cell dissociation was confirmed with a microscope. The dissociation was stopped with 1 ml ice cold PBS supplemented with 10% fetal bovine serum (FBS). Afterwards the cells were immediately pelleted at 300× *g* for 5 min at 4°C. Subsequently, the supernatant was discarded and cells were suspended in 2 ml PBS/10%FBS and pelleted again at 300× *g* for min at 4°C. Then cells were suspended in PBS/0.01%BSA and pelleted again (300× g for 5 min at 4°C), suspended in 1 ml PBS/0.01%BSA, and passed through a 30 μM filter mesh (Miltenyi MACS smart strainer). Viability was then investigated with the Trypan-blue exclusion test and cell concentration was determined with a hemocytometer and adjusted to 200,000 cells/ml for Drop-seq.

**Drop-seq** Uniformly dispersed 1 nl-sized droplets were generated using self-built polydimethylsiloxane (PDMS) microfluidic co-flow devices on the basis of the AutoCAD design provided by the McCarroll group. The devices were treated with a water repellant solution (Aquapel) to create a hydrophobic channel surface. Drop-Seq runs followed closely the procedure published by Macosko et al. (Online Dropseq protocol v. 3.1 http://mccarrolllab.com/dropseq/). Barcoded beads (ChemGenes Corp., Wilmington, MA), suspended in lysis buffer, were co-flown with a single cell suspension and a droplet generation mineral oil (QX200, Bio-Rad Laboratories). Resulting droplets were collected in a 50 ml tube and immediately disrupted after adding 30 ml high-salt saline-sodium citrate buffer (6×SSC) and 1 ml perfluorooctanol. Subsequently, captured mRNA’s were reverse transcribed for 2 hours using 2,000 U of the Maxima H Minus Reverse Transcriptase (ThermoFisher) followed by an exonuclease treatment for 45 minutes to remove unextended primers. After two washing steps with 6×SSC buffer about 70,000 remaining beads (60% of input beads) were aliquoted (5,000 beads per 50 μl reaction) and PCR-amplified (5 cycles at 65°C and 12 cycles at 67°C annealing temperature). Aliquots of each PCR reaction were pooled and double-purified using 0.5× volume of Agencourt AMPure XP beads (# A63881, Beckman Coulter) and finally eluted in 10 μl EB buffer. Quality and quantity of the amplified cDNAs were analyzed on a BioAnalyzer High Sensitivity DNA Chip (Agilent Technologies, Santa Clara, CA). About 600 pg cDNA was fragmented and amplified (17 cycles) to generate a next-generation sequencing library by using the Nextera XT DNA sample preparation kit (Illumina). The libraries were purified, quantified (Agilent High sensitivity DNA chip), and then sequenced (paired end 26×115 bases) on the Illumina HiSeq2500 platform. Custom primer (5’-GCCTGTCCGCGGAAGCAGTGGTATCAACGCAGAGTAC-3’) was used for the first sequence read to identify all different cell barcodes und unique molecular identifier (UMI) sequences.

**Bioinformatics analysis:** Studies have reported that low-coverage sequencing of single cells (~10,000 reads) is sufficient for unbiased classification of diverse cell types in heterogeneous tissue; finer sub-clusters requires moderately higher depths (~50,000 reads)(Pollen et al., 2014),(Bacher and Kendziorski, 2016),(Ntranos et al., 2016). The quality of the fastq files from the sequencer were first checked using FastQC (v0.11.4) and trimming of the 3’reads were done in case of base sequence quality < 30. Next, using the tools embedded in Picard tools (picard-tools-1.115) and the DropSeq analysis pipeline developed by the McCarroll lab (http://mccarrolllab.com/dropseq/), the fastq files were processed and the data matrix table containing the gene expression of the barcoded cells was generated. Individual cells were labeled with barcodes, and transcripts within each cell were tagged with distinct UMIs (Unique Molecular Identifiers) in order to determine absolute transcript abundance. Based on the total number reads per barcode a cumulative distribution plot was generated; the deflection point in the plot indicates the number of barcoded cells to be used for further analysis. We combined the expression from three fetal kidney datasets for this study. Combination of statistical R packages (Diplyr, Matrix, Seurat, and SVA) was used for our downstream analyses. We removed from our analysis all cells with < 500 genes and also the cells in which the mitochondrial transcripts constituted more than 10% of the total transcripts counts. Transcript counts within each cell (each column in the data matrix) were then median normalized and the data matrix was batch corrected using Combat (Shekhar et al., 2016). Next, we identified the highly variable genes across the single cells and performed principal component analysis using these genes. Significant PCs were then used to for an unsupervised clustering of the cells. Differentially expressed cell-type markers for each cluster were determined using the embedded tobit model in the Seurat R package. The clusters were viewed in two-dimensional frame using tSNE clustering (Shekhar et al., 2016).

### Trajectory estimation from single-cell RNA-seq data

Single-cell RNA-seq data was normalized and log-transformed by R package Seurat. The transformed data was filtered by the following criteria: 1. Include only genes with detected expression in at least 3 cells. 2. Include only cells with at least 1000 genes detected and with less than 5% of total reads being mitochondria reads.

To remove technical variations, we regressed out number of genes expressed, percentage of mitochondria reads, and batch variables via a linear model. To control for confounding effects of cell cycle to trajectory estimation, we subtracted gene expression level of non-cell cycle genes by a baseline estimated using expression of only cell cycle genes. Specifically, we used a curated cell cycle gene list of 1946 genes from Barron and Li(Barron and Li, 2016).; for each non-cell cycle gene, we fitted a ridge regression model to predict its log-transformed expression level from cell-cycle genes, using cv.glmnet function in R package glmnet. The regularization parameter was automatically selected based on 10-fold cross-validation. With the fitted models we computed the residuals of each non-cell cycle gene and use the residuals as input for trajectory analysis.

Trajectory analysis was performed with a method described in Zhou and Troyanskaya (manuscript in preparation). To identify genes with significant expression change along the trajectory, for each trajectory segment of interest, we tested the significance of each gene by fitting a generalized additive model(GAM) with thin plate regression spline to predict expression level from the cell order in the trajectory segment using the gam function of R package mgcv. The significance level of the gene is the significance level of cell order in the GAM model.

### Immunofluorescence

Kidneys were fixed overnight in 4%PFA and processed for frozen sectioning and incubation as previously reported (Cebrian, 2012). Thin 5-micron sections were incubated at 4C overnight with primary antibodies against the following epitopes: Decorin (R&D systems, AF143), ETV4 (Proteintech, 10684-1-AP), ACTA2 (Sigma, C6198), PAX8 (Proteintech, 10336-1-AP), RET (R&D, AF1485), Synaptopodin (Proteintech, 21064-1-AP) and TenascinC (R&D, MAB2138). Alexa 488-conjugated secondary antibodies were purchased from Jackson ImmunoResearch and incubated at 37C for 3h. Imaging was performed on an Olympus IX71 inverted microscope or a Nikon A1 confocal microscope.

## Acknowledgements

We are indebted to Steve Potter for sharing the cold protease protocol before publication.

## Data availability

Datasets generated in this study are available from GEO repository: https://www.ncbi.nlm.nih.gov/geo/query/acc.cgi?acc=GSE109205

---

Supplementary figure 1: A: Heatmap identifying 19 clusters. B: Correlation plot for clusters 3, 4, 16 and 18.

Supplementary figure 2: t-SNE cluster map of the expression levels for EYA1, TMEM100, ELAVL4, ECEL1, COL2A1 and ETV1. Each dot represents an individual cell and color intensity reflects gene expression levels.

Supplementary figure 3: t-SNE cluster map of the expression levels for WNT4, WNT7B, WNT5A and WNT11. Each dot represents an individual cell and color intensity reflects gene expression levels.

Supplementary figure 4: t-SNE cluster map of the expression levels for FGFR1, FGFR2, FGF7 and FGF8. Each dot represents an individual cell and color intensity reflects gene expression levels.

Supplementary table 1: List of genes differentially expressed by cells within each cluster. Pct1: Percentage of cells in the cluster of interest. Pct2: Percentage of cells in all other clusters.

## References

Adam, M., Potter, A.S., Potter, S.S., 2017. Psychrophilic proteases dramatically reduce singlecell RNA-seq artifacts: a molecular atlas of kidney development. Development 144, 3625–3632.

Bacher, R., Kendziorski, C., 2016. Design and computational analysis of single-cell RNA-sequencing experiments. Genome Biology 17, 63.

Barak, H., Huh, S.H., Chen, S., Jeanpierre, C., Martinovic, J., Parisot, M., Bole-Feysot, C., Nitschke, P., Salomon, R., Antignac, C., Ornitz, D.M., Kopan, R., 2012. FGF9 and FGF20 maintain the stemness of nephron progenitors in mice and man. Dev Cell 22, 1191–1207.

Barron, M., Li, J., 2016. Identifying and removing the cell-cycle effect from single-cell RNA-Sequencing data. Sci Rep 6, 33892.

Boyle, S.C., Kim, M., Valerius, M.T., McMahon, A.P., Kopan, R., 2011. Notch pathway activation can replace the requirement for Wnt4 and Wnt9b in mesenchymal-to-epithelial transition of nephron stem cells. Development 138, 4245–4254.

Boyle, S.C., Liu, Z., Kopan, R., 2014. Notch signaling is required for the formation of mesangial cells from a stromal mesenchyme precursor during kidney development. Development 141, 346–354.

Cacalano, G., Farinas, I., Wang, L.C., Hagler, K., Forgie, A., Moore, M., Armanini, M., Phillips, H., Ryan, A.M., Reichardt, L.F., Hynes, M., Davies, A., Rosenthal, A., 1998. GFRalpha1 is an essential receptor component for GDNF in the developing nervous system and kidney. Neuron 21, 53–62.

Carroll, T.J., Park, J.S., Hayashi, S., Majumdar, A., McMahon, A.P., 2005. Wnt9b plays a central role in the regulation of mesenchymal to epithelial transitions underlying organogenesis of the mammalian urogenital system. Dev Cell 9, 283–292.

Cebrian, C., 2012. Fluorescent immunolabeling of embryonic kidney samples. Methods Mol Biol 886, 251–259.

Cebrian, C., Asai, N., D'Agati, V., Costantini, F., 2014. The number of fetal nephron progenitor cells limits ureteric branching and adult nephron endowment. Cell reports 7, 127–137.

Cheng, H.T., Kim, M., Valerius, M.T., Surendran, K., Schuster-Gossler, K., Gossler, A., McMahon, A.P., Kopan, R., 2007. Notch2, but not Notch1, is required for proximal fate acquisition in the mammalian nephron. Development 134, 801–811.

Chi, X., Michos, O., Shakya, R., Riccio, P., Enomoto, H., Licht, J.D., Asai, N., Takahashi, M., Ohgami, N., Kato, M., Mendelsohn, C., Costantini, F., 2009. Ret-dependent cell rearrangements in the Wolffian duct epithelium initiate ureteric bud morphogenesis. Dev Cell 17, 199–209.

Das, A., Tanigawa, S., Karner, C.M., Xin, M., Lum, L., Chen, C., Olson, E.N., Perantoni, A.O., Carroll, T.J., 2013. Stromal-epithelial crosstalk regulates kidney progenitor cell differentiation. Nat Cell Biol 15, 1035–1044.

Ehrenfels, C.W., Carmillo, P.J., Orozco, O., Cate, R.L., Sanicola, M., 1999. Perturbation of RET signaling in the embryonic kidney. Dev Genet 24, 263–272.

Garrod, D.R., Fleming, S., 1990. Early expression of desmosomal components during kidney tubule morphogenesis in human and murine embryos. Development 108, 313–321.

Golden, J.P., DeMaro, J.A., Osborne, P.A., Milbrandt, J., Johnson, E.M., Jr., 1999. Expression of neurturin, GDNF, and GDNF family-receptor mRNA in the developing and mature mouse. Exp Neurol 158, 504–528.

Grieshammer, U., Cebrian, C., Ilagan, R., Meyers, E., Herzlinger, D., Martin, G.R., 2005. FGF8 is required for cell survival at distinct stages of nephrogenesis and for regulation of gene expression in nascent nephrons. Development 132, 3847–3857.

Grobstein, C., 1953. Inductive epitheliomesenchymal interaction in cultured organ rudiments of the mouse. Science 118, 52–55.

Halt, K.J., Parssinen, H.E., Junttila, S.M., Saarela, U., Sims-Lucas, S., Koivunen, P., Myllyharju, J., Quaggin, S., Skovorodkin, I.N., Vainio, S.J., 2016. CD146(+) cells are essential for kidney vasculature development. Kidney Int 90, 311–324.

Hillier, B.J., Vacquier, V.D., 2003. Amassin, an olfactomedin protein, mediates the massive intercellular adhesion of sea urchin coelomocytes. J Cell Biol 160, 597–604.

Huang, J., Arsenault, M., Kann, M., Lopez-Mendez, C., Saleh, M., Wadowska, D., Taglienti, M., Ho, J., Miao, Y., Sims, D., Spears, J., Lopez, A., Wright, G., Hartwig, S., 2013. The transcription factor Sry-related HMG box-4 (SOX4) is required for normal renal development in vivo. Dev Dyn 242, 790–799.

Huang, L., Xiao, A., Choi, S.Y., Kan, Q., Zhou, W., Chacon-Heszele, M.F., Ryu, Y.K., McKenna, S.,Zuo, X., Kuruvilla, R., Lipschutz, J.H., 2014. Wnt5a is necessary for normal kidney development in zebrafish and mice. Nephron Exp Nephrol 128, 80–88.

Ihara, K., Asanuma, K., Fukuda, T., Ohwada, S., Yoshida, M., Nishimori, K., 2014. MAGI-2 is critical for the formation and maintenance of the glomerular filtration barrier in mouse kidney. Am J Pathol 184, 2699–2708.

Itoh, M., Nakadate, K., Horibata, Y., Matsusaka, T., Xu, J., Hunziker, W., Sugimoto, H., 2014. The structural and functional organization of the podocyte filtration slits is regulated by Tjp1/ZO-1 PLoS One 9, e106621.

Kazama, I., Mahoney, Z., Miner, J.H., Graf, D., Economides, A.N., Kreidberg, J.A., 2008. Podocyte-derived BMP7 is critical for nephron development. J Am Soc Nephrol 19, 2181–2191.

Kobayashi, A., Valerius, M.T., Mugford, J.W., Carroll, T.J., Self, M., Oliver, G., McMahon, A.P., 2008. Six2 defines and regulates a multipotent self-renewing nephron progenitor population throughout mammalian kidney development. Cell Stem Cell 3, 169–181.

Koehler, S., Tellkamp, F., Niessen, C.M., Bloch, W., Kerjaschki, D., Schermer, B., Benzing, T., Brinkkoetter, P.T., 2016. Par3A is dispensable for the function of the glomerular filtration barrier of the kidney. Am J Physiol Renal Physiol 311, F112–119.

Koike, H., Zhang, R.R., Ueno, Y., Sekine, K., Zheng, Y.W., Takebe, T., Taniguchi, H., 2017. Nutritional modulation of mouse and human liver bud growth through a branched-chain amino acid metabolism. Development 144, 1018–1024.

Kuure, S., Chi, X., Lu, B., Costantini, F., 2010. The transcription factors Etv4 and Etv5 mediate formation of the ureteric bud tip domain during kidney development. Development 137, 1975–1979.

Lai Wing Sun, K., Correia, J.P., Kennedy, T.E., 2011. Netrins: versatile extracellular cues with diverse functions. Development 138, 2153–2169.

Lindenmeyer, M.T., Eichinger, F., Sen, K., Anders, H.J., Edenhofer, I., Mattinzoli, D., Kretzler, M., Rastaldi, M.P., Cohen, C.D., 2010. Systematic analysis of a novel human renal glomerulus-enriched gene expression dataset. PLoS One 5, e11545.

Lu, B.C., Cebrian, C., Chi, X., Kuure, S., Kuo, R., Bates, C.M., Arber, S., Hassell, J., MacNeil,L., Hoshi, M., Jain, S., Asai, N., Takahashi, M., Schmidt-Ott, K.M., Barasch, J., D'Agati, V., Costantini, F., 2009. Etv4 and Etv5 are required downstream of GDNF and Ret for kidney branching morphogenesis. Nat Genet 41, 1295–1302.

Lu, Y., Ye, Y., Bao, W., Yang, Q., Wang, J., Liu, Z., Shi, S., 2017. Genome-wide identification of genes essential for podocyte cytoskeletons based on single-cell RNA sequencing. Kidney Int.

Mason, I.J., Fuller-Pace, F., Smith, R., Dickson, C., 1994. FGF-7 (keratinocyte growth factor) expression during mouse development suggests roles in myogenesis, forebrain regionalisation and epithelial-mesenchymal interactions. Mechanisms of development 45, 15–30.

Matsumoto, S., Fujii, S., Sato, A., Ibuka, S., Kagawa, Y., Ishii, M., Kikuchi, A., 2014. A combination of Wnt and growth factor signaling induces Arl4c expression to form epithelial tubular structures. The EMBO journal 33, 702–718.

Meerabux, J.M., Ohba, H., Fukasawa, M., Suto, Y., Aoki-Suzuki, M., Nakashiba, T., Nishimura,S.,Itohara, S., Yoshikawa, T., 2005. Human netrin-G1 isoforms show evidence of differential expression. Genomics 86, 112–116.

Mugford, J.W., Sipila, P., McMahon, J.A., McMahon, A.P., 2008. Osr1 expression demarcates a multi-potent population of intermediate mesoderm that undergoes progressive restriction to an Osr1-dependent nephron progenitor compartment within the mammalian kidney. Dev Biol 324, 88–98.

Muller, U., Wang, D., Denda, S., Meneses, J.J., Pedersen, R.A., Reichardt, L.F., 1997. Integrin alpha8beta1 is critically important for epithelial-mesenchymal interactions during kidney morphogenesis. Cell 88, 603–613.

Ng, L.J., Tam, P.P., Cheah, K.S., 1993. Preferential expression of alternatively spliced mRNAs encoding type II procollagen with a cysteine-rich amino-propeptide in differentiating cartilage and nonchondrogenic tissues during early mouse development. Dev Biol 159, 403–417.

Ntranos, V., Kamath, G.M., Zhang, J.M., Pachter, L., Tse, D.N., 2016. Fast and accurate single-cell RNA-seq analysis by clustering of transcript-compatibility counts. Genome Biology 17, 112.

O'Brien, L.L., Guo, Q., Lee, Y., Tran, T., Benazet, J.D., Whitney, P.H., Valouev, A., McMahon, A.P., 2016. Differential regulation of mouse and human nephron progenitors by the Six family of transcriptional regulators. Development 143, 595–608.

Ohse, T., Chang, A.M., Pippin, J.W., Jarad, G., Hudkins, K.L., Alpers, C.E., Miner, J.H., Shankland, S.J., 2009. A new function for parietal epithelial cells: a second glomerular barrier. Am J Physiol Renal Physiol 297, F1566–1574.

Pachnis, V., Mankoo, B., Costantini, F., 1993. Expression of the c-ret proto-oncogene during mouse embryogenesis. Development 119, 1005–1017.

Park, J.S., Valerius, M.T., McMahon, A.P., 2007. Wnt/beta-catenin signaling regulates nephron induction during mouse kidney development. Development 134, 2533–2539.

Patrakka, J., Xiao, Z., Nukui, M., Takemoto, M., He, L., Oddsson, A., Perisic, L., Kaukinen, A., Szigyarto, C.A., Uhlen, M., Jalanko, H., Betsholtz, C., Tryggvason, K., 2007. Expression and subcellular distribution of novel glomerulus-associated proteins dendrin, ehd3, sh2d4a, plekhh2, and 2310066E14Rik. J Am Soc Nephrol 18, 689–697.

Perantoni, A.O., Timofeeva, O., Naillat, F., Richman, C., Pajni-Underwood, S., Wilson, C., Vainio, S., Dove, L.F., Lewandoski, M., 2005. Inactivation of FGF8 in early mesoderm reveals an essential role in kidney development. Development 132, 3859–3871.

Pichel, J.G., Shen, L., Sheng, H.Z., Granholm, A.C., Drago, J., Grinberg, A., Lee, E.J., Huang,S.P., Saarma, M., Hoffer, B.J., Sariola, H., Westphal, H., 1996. Defects in enteric innervation and kidney development in mice lacking GDNF. Nature 382, 73–76.

Pietila, I., Prunskaite-Hyyrylainen, R., Kaisto, S., Tika, E., van Eerde, A.M., Salo, A.M., Garma, L., Miinalainen, I., Feitz, W.F., Bongers, E.M., Juffer, A., Knoers, N.V., Renkema, K.Y., Myllyharju, J., Vainio, S.J., 2016. Wnt5a Deficiency Leads to Anomalies in Ureteric Tree Development, Tubular Epithelial Cell Organization and Basement Membrane Integrity Pointing to a Role in Kidney Collecting Duct Patterning. PLoS One 11, e0147171.

Pollen, A.A., Nowakowski, T.J., Shuga, J., Wang, X., Leyrat, A.A., Lui, J.H., Li, N., Szpankowski, L., Fowler, B., Chen, P., Ramalingam, N., Sun, G., Thu, M., Norris, M., Lebofsky, R., Toppani, D., Kemp, D., Wong, M., Clerkson, B., Jones, B.N., Wu, S., Knutsson, L., Alvarado, B., Wang, J., Weaver, L.S., May, A.P., Jones, R.C., Unger, M.A., Kriegstein, A.R., West, J.A.A., 2014. Low-coverage single-cell mRNA sequencing reveals cellular heterogeneity and activated signaling pathways in developing cerebral cortex. Nature biotechnology 32, 1053–1058.

Riccio, P., Cebrian, C., Zong, H., Hippenmeyer, S., Costantini, F., 2016. Ret and Etv4 Promote Directed Movements of Progenitor Cells during Renal Branching Morphogenesis. PLoS biology 14, e1002382.

Rumballe, B.A., Georgas, K.M., Combes, A.N., Ju, A.L., Gilbert, T., Little, M.H., 2011. Nephron formation adopts a novel spatial topology at cessation of nephrogenesis. Dev Biol 360, 110–122.

Schuchardt, A., D'Agati, V., Larsson-Blomberg, L., Costantini, F., Pachnis, V., 1994. Defects in the kidney and enteric nervous system of mice lacking the tyrosine kinase receptor Ret. Nature 367, 380–383.

Shakya, R., Watanabe, T., Costantini, F., 2005. The role of GDNF/Ret signaling in ureteric bud cell fate and branching morphogenesis. Dev Cell 8, 65–74.

Shekhar, K., Lapan, S.W., Whitney, I.E., Tran, N.M., Macosko, E.Z., Kowalczyk, M., Adiconis, X., Levin, J.Z., Nemesh, J., Goldman, M., McCarroll, S.A., Cepko, C.L., Regev, A., Sanes, J.R., 2016. Comprehensive Classification of Retinal Bipolar Neurons by Single-Cell Transcriptomics. Cell 166, 1308–1323. e1330.

Sistani, L., Duner, F., Udumala, S., Hultenby, K., Uhlen, M., Betsholtz, C., Tryggvason, K., Wernerson, A., Patrakka, J., 2011. Pdlim2 is a novel actin-regulating protein of podocyte foot processes. Kidney Int 80, 1045–1054.

Surendran, K., Boyle, S., Barak, H., Kim, M., Stomberski, C., McCright, B., Kopan, R., 2010. The contribution of Notch1 to nephron segmentation in the developing kidney is revealed in a sensitized Notch2 background and can be augmented by reducing Mint dosage. Dev Biol 337, 386–395.

Tsukahara, Y., Tanaka, M., Miyajima, A., 2011. TROP2 expressed in the trunk of the ureteric duct regulates branching morphogenesis during kidney development. PLoS One 6, e28607.

Wada, J., Kumar, A., Ota, K., Wallner, E.I., Batlle, D.C., Kanwar, Y.S., 1997. Representational difference analysis of cDNA of genes expressed in embryonic kidney. Kidney Int 51, 1629–1638.

Xu, P.X., Adams, J., Peters, H., Brown, M.C., Heaney, S., Maas, R., 1999. Eya1-deficient mice lack ears and kidneys and show abnormal apoptosis of organ primordia. Nat Genet 23, 113–117.

Yun, K., Perantoni, A.O., 2017. Hydronephrosis in the Wnt5a-ablated kidney is caused by an abnormal ureter-bladder connection. Differentiation 94, 1–7.

Zhang, S., Lin, Y., Itaranta, P., Yagi, A., Vainio, S., 2001. Expression of Sprouty genes 1, 2 and 4 during mouse organogenesis. Mechanisms of development 109, 367–370.

Zhao, H., Kegg, H., Grady, S., Truong, H.T., Robinson, M.L., Baum, M., Bates, C.M., 2004. Role of fibroblast growth factor receptors 1 and 2 in the ureteric bud. Dev Biol 276, 403–415.

